# Design and strain selection criteria for bacterial communication networks

**DOI:** 10.1101/013383

**Authors:** Claudio Angione, Giovanni Carapezza, Jole Costanza, Pietro Lió, Giuseppe Nicosia

## Abstract

In this paper we discuss data and methodological challenges for building bacterial communication networks using two examples: *Escherichia coli* as a flagellate bacterium and of *Geobacter sulfurreducens* as a biofilm forming bacterium. We first highlight the link between the bacterial network communication design with respect to metabolic information processing design. The potentialities of designing routing network schemes described previously in literature and based on bacteria motility and genetic message exchanges will depend on the genes coding for the intracellular and intercellular signalling pathways. In bacteria, the “mobilome” is related to horizontal gene transfer. Bacteria trade off the acquisition of new genes which could improve their survival (and often their communication bandwidth), keeping their genome enough small to assure quick DNA replication and increase fast the biomass to speed up cell division. First, by using a multi-objective optimisation procedure, we search for the optimal trade off between energy production, which is a requirement for the motility, and the biomass growth, which is related to the overall survival and fitness of the bacterium. We use flux balance analysis of genome-scale biochemical network of *Escherichia coli* k-13 MG1655. Then, as a second case study we analyze the electric properties and biomass trade-off of the bacterium *Geobacter sulfurreducens* which constructs an electric biofilm where electrons move across the nanowires. Here we discuss the potentialities of optimisation methodologies to design and select bacterial strains with desiderata properties. The optimisation methodologies establish also a relation between metabolism, network communication and computing. Moreover, we point to genetic design and synthetic biology as key areas to develop bacterial nano communication networks.

## Introduction: Communications in bacteria

Bacteria colonise every environment; for example the human intestinal microbiota may contain 10^13^ to 10^14^ bacteria whose overall gene diversity (“microbiome”) contains at least 100 times as many genes as our own. Our microbiome has a significantly enriched metabolism of glycans, amino acids, and pathway-mediated biosynthesis of vitamins and isoprenoids. The study of metabolism is the key aspect to evaluate the role of bacteria in the environments, as well as to design new biotechnologically useful strains. In general, the study of metabolism can provide key insights into the understanding of the overall behaviour of bacteria and, importantly, into their phenotype-genotype relationships. The experimental investigation of the metabolic capabilities of a given organism is highly demanding in terms of both costs and wet-lab resources. Conversely, an in silico analysis of the overall metabolic capabilities of the system can help in predicting or designing phenotypes or, at least, reduce the choice of useful experiments.

Bacteria could be used to build nano communication networks that operate in microfluidic devices, body area networks or other environments [1]. Bacteria are constrained by an impermeable membrane. They have a system of surface proteins regulating the exchange of information, and a molecular system inside the cell that interprets the external information and acts upon. The fitness of a bacterium is particularly concentrated on the speed of dividing, but the cell division time depends on reaching a certain biomass. In order to achieve a biomass, the bacterium needs energy to locate a source of food and move towards it. A bacterium typically swims by alternating straight runs with short periods of tumbles that randomly reorientate the next run. Motile bacteria suppress tumbles when they head either up concentration gradients of attractants or down gradients of repellents. Motile bacteria synthesise proteins for chemotaxis including flagella formation when the substrate concentration, i.e. food, becomes low. The synthesis and function of the flagellar and chemotaxis system requires the expression of a network of more than 50 genes, therefore it is genomically and metabolic expensive. Using the proteins coded by those genes, a bacterium uses receptors to sense the spatial gradient and compares the instantaneous concentration of carbon sources. Although existing models of bacterial chemotaxis do not take into account the tight coupling with the metabolism, it is known that the metabolism modulates chemotaxis and motility behaviour. In this paper, we study the metabolism as a trade -off between energy (required for motility) and the biomass (required for the growth). A decrease in biomass due to starvation would require spending resources towards searching new source of food and therefore accomplishing chemotaxis specific signal transduction, through the direct modulation of flagellar rotation.

In the next sections, we describe the methodology for designing strains (Pareto fronts and flux balance analysis (FBA)), the additional information from other omics we may use, the relations between metabolism, computing and communications (see also [2], [3]).

## I. Flux balance analysis

The flux balance analysis (FBA) approach provides the solutions (reaction fluxes) satisfying the optimisation of the most efficient use of available resources, such as nutrients [4]. Organic compounds are converted to carbon skeletons for the synthesis of various cell components and for the production of energy. This is possible through the regulation of the reactions of anabolism and catabolism. We believe that the bacterium behaviour could be analysed using multi-optimisation techniques [5]. The result of the multi-objective optimisation is not a single solution (such as in a single optimisation problem), but a set of non-dominated points, which constitutes the Pareto surface. Therefore, we can seek the best trade-off design for the set of pathways considered, leading to optimise simultaneously multiple cellular functions of interest. For each Pareto optimal solution, it will be possible to compute the robustness, the sensitivity and the identifiability integrated with all the available data. The information on the sensitivity (the elements that have a large influence on the system outputs are considered sensitive) of biological networks and pathways could be used to suggest where and how much to modify a metabolic network.

A three-dimensional Pareto front can help reaching the trade-off among biomass production, genome size and the metabolism associated with them, and energy for chemotaxis and movement (Figure 1). For example bacterial needs to increase the biomass (biomass size influences cell division), keep low the genome size in order to replicate quickly. Probably for this reason, genes transitorily useful are kept on accessory chromosomes which could be replicated in parallel and be lost or kept. We assume that the energy for the location of food source is in high demand and has higher priority than the biomass growth and division. It is noteworthy that metabolic and genome design could produce bacteria with different motility and different tumble frequencies to fit ad hoc network topologies. Furthermore, the capacity to respond to gradients could be tuned by expressing different types and amount of receptor proteins or sensor pathways.

**Fig. 1.**
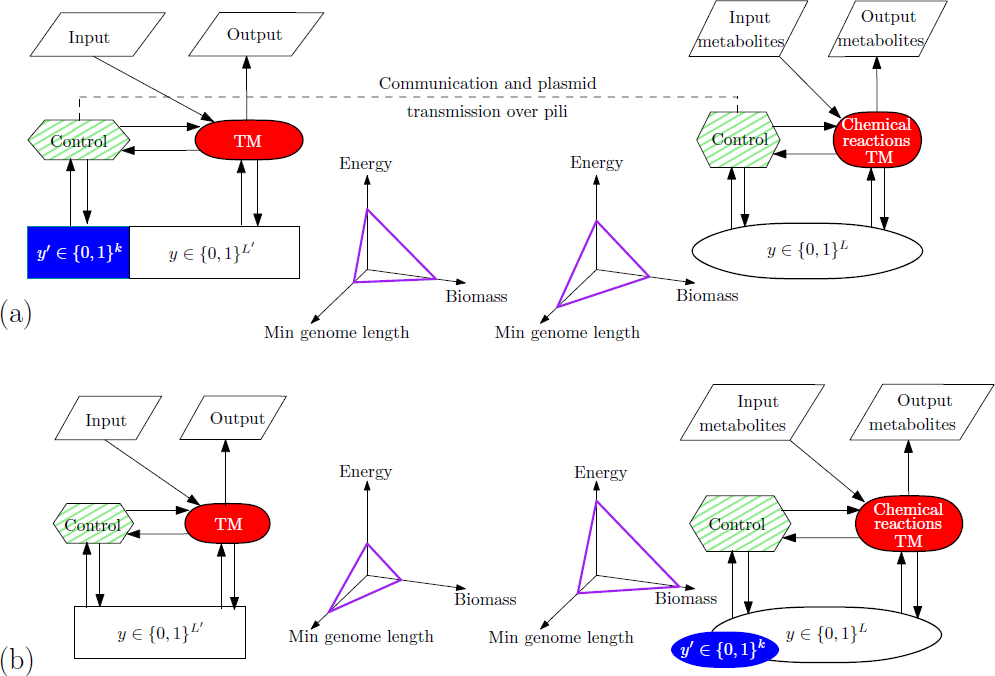
Trade-off among biomass production (which affects the cell duplication time), the minimal size of the genome (which affects the genome replication time), and energy for chemotaxis, movement and other cell activities. In a first step (a), the bacterium on the left sends a piece of genome to the bacterium on the right, which engulfs the new piece of code enriching its genome (b). This allows to produce more enrgy for movement and biomass, but at the cost of an increased genome size.

**Fig. 2.**
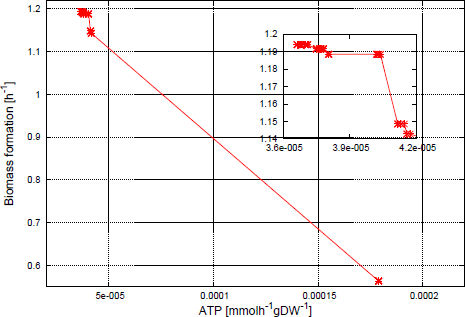
Pareto front of the biomass (*y*-axis) versus energy (ATP) (*x*-axis) in *E. coli;* a more “energetic” strain would represent a choice towards an increased motility; the biomass choice would represent a choice towards a faster replicating strain.

### Designing properties using Pareto front guidance

When a system cannot optimise all the tasks it performs, a trade-off between contrasting objectives can be obtained using a multi-objective optimisation technique For example, two communicating organisms can harness the many-objective Pareto optimality to find a trade-off decision that allows to define their behaviour. The Pareto front allows to maximise or minimise two or more target metabolites in an organism, thus obtaining new optimal strains specialised in many aims concurrently. By adopting a trade-off strategy, an organism is able to optimise simultaneously several biotechnological targets, e.g. the input and the output of the computation it carries out. Given *r* objective functions *f*_1_, …, *f*_*r*_ to optimise, the problem of optimising in a multi-objective fashion can be formalised as

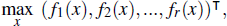

where *x* is the variable in the search space. Without loss of generality, in the definition all the functions are maximised (however, minimising a function *f*_*i*_ is equivalent to maximising −*f*_*i*_). The output of a multi-objective routine is a set of Pareto optimal points, which constitute the *Pareto front*. A solution *y*\* is Pareto optimal if there does not exist a point *y* such that *f*(*y*) dominates *f*(*y*\*). Formally, *y*\* is Pareto optimal if ∄ *y* s.t. *f*_*i*_(*y*) > *f*_*i*_(*y*\*) ∀*i* = 1, …, *r*, where *f* is the vector of *r* objective functions that have to be maximised in the objective space.

Bacteria can be genetically modified and the changes described in the Pareto-front framework to find the best tradeoff between two or more requirements. Since the bacterium has always more than one functions, the decision whether to communicate with another bacterium has to take into account the output of an internal multi-objective optimisation routine. For instance, an *E. coli* whose objectives are the production of acetate and biomass, obtains the Pareto front in Figure 4. (the model taken into account is the *E. coli* by orth et al. [6].) A Pareto front produced by an organism is the set of all the phenotypes that overcome all the feasible phenotypes dominated on all tasks [7]. Communicating with other bacteria is intended to increase the computational capability of the bacterium, and therefore moves the Pareto front towards the best unfeasible point, which is located at the top right of the acetate-biomass graph. Nevertheless, this can decrease the capability of the bacterium to produce a third-objective, and therefore the decision may require a three-objective optimisation routine. The communication happens when two bacteria share DNA fragments (see Figure 6).

**Fig. 4.**
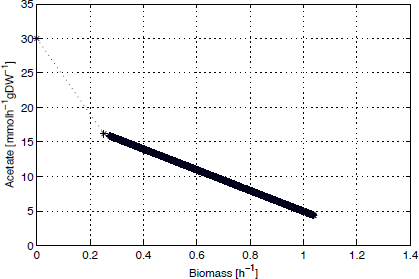
Result of the two-objective optimisation routine carried out on the *E. coli* model. Since the communication among bacteria allows to share DNA fragments, ant therefore increases their computational capabilities towards one or more objectives (e.g., acetate and biomass) in which a bacterium specialises.

**Fig. 5.**
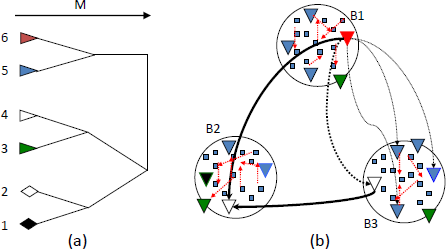
The tree in (a) describes the duplication events, or LGT. Recent events are separated by short distance in the *x* axis, i.e. events with a low number of mutations occurred. In (b), the LGT events occurring in a network of three bacteria (B1-3); in green are the sensor or flagellar proteins; in red the protein interactions. The LGT events generate often similar but not equal metabolic networks.

**Fig. 6.**
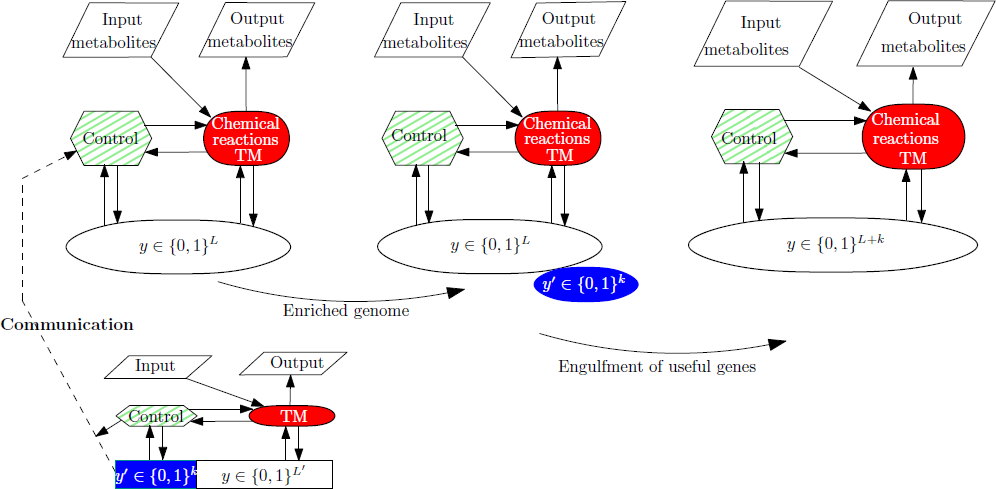
Communication between two bacteria. Communication between bacteria (left) allows the small bacterium to copy and send a piece of genome to the larger one, which engulfs the new piece of code and increases its computational capability (centre). Therefore, the genome is enriched with respect to the objectives that the maximises (right).

### The mobilome: conjugation for network communication

Bacterial conjugation involves DNA transfer between donor and recipient called *lateral gene transfer* (LGT), which has great importance in building the genomes of prokaryotic organisms. The LGT from bacteria to humans is more likely to occur in tumour samples than in healthy somatic cells [8], [9].

In bacteria, horizontal gene transfer is often mediated by conjugative genetic sequences that transfer directly from cell to cell. Integrative and conjugative sequences are mobile genetic elements that reside within a host genome but can excise to form a circle and transfer by conjugation to recipient cells. The LGT device in the cell is assembled each time there are conditions for exchange. Gene duplication has long been recognised as an important mechanism for the creation of new gene functions [10]. Note that due to LGT half of all *S. aureus* infections in the world are resistant to penicillin, methicillin, tetracycline and erythromycin. Vancomycin resistant bacteria were first identified in Japan in 1996, and strains have since been found in hospitals in England, France and the US. LGT facilitates efficient communication among the cells, and provides them access to a global prokaryote genome (superorganism), preventing massive extinctions of genes. Therefore, FBA could be meaningful to provide a methodological means to integrate superorganism metabolism. In other words, FBA could model steady state conditions for both metabolic, single and multiple compartment systems and ecosystems (natural such as free external environments, gut and other organs or endosymbiontic environments and artificial such as microfluidic devices or other devices). The acquisition of novel useful information comes at the cost of replicating a larger genome, spending energy content molecules or intermediate of reactions not always directly related to cell growth or division, increasing the resources for control. The competition among bacteria favours efficient resource (number of genes) management, and therefore redundant or rarely used information is frequently lost. This results in a maintenance of the genome size for the majority of bacterial species which could keep additional information (such as genes coding for proteins important for survival in antibiotics or metal rich environment) on plasmids (accessory chromosomes). We may think that the trade off between genome size and metabolic richness could be described by a Pareto front representing the two-objective optimisation.

## II. Targets for designing bacterial communication

The surface of the bacteria contains pili, which allows the bacteria to come together and form contacts. The pili are 1 *μ*m long and 6-7 nm diameter; they have extension speed of 400*nm*/*s* and 6*nm*/*s*. In many cases bacteria form colonies and biofilms which make the encounters highly probable. Then gene transfer often happens. The relationship between time and amount of transferred genetic information is linear (after a certain delay due to the assembling of the protein machinery) and accurate enough to be used for identifying the order of the genes through so called interrupted-mating. The transfer process takes several minutes to start due to the assembling of the related structures, but proceeds quickly. For example, a sequence of one hundred bases could take 5 minutes for the transfer, while 5 million bases will take approximately 100 minutes. The important property of conjugation is the meeting distance between the bacteria.

Conjugation is mediated by self-transmissible plasmids such as F-Plasmids, as well as phage-like sequences that have been integrated into the bacterial chromosome, such as Integrative and Conjugative Elements (ICEs). Both conjugative plasmids and ICEs can mediate the transfer of mobile elements by sharing their conjugative machinery. An important observation is that many bacteria grow in chains, i.e. dense communities of cells, where the presence of conjugative elements in cells can contribute to the formation of such communities.

When acquired by one cell in a chain, ICEs spread rapidly from cell to cell within the chain by additional sequential conjugation events. This intra-chain conjugation is inherently more efficient than conjugation that is due to chance encounters between individual cells. Therefore, although the process is slow because it requires building a protein complex, it can quickly spread, where a single donor cell can convert a population of recipient cells to donor cell status via a process similar to epidemic spreading. Conjugation requires coupling proteins that links the transferosome (*a* type IV secretion system in Gram-negative bacteria) to the relaxosome, a nucleoprotein complex at the origin of transfer. Although broadly speaking, the transfer systems appear to be able to drill a hole through any recipient cell envelope and start the DNA transfer in a recipient-independent manner, there are several factors and security check that affect the process. The transfer potential of these transfer regions depends on the integration of many signals in response to environmental and physiological cues. Conjugative elements can be narrow or broad host range, depending on their ability to be established and maintained in the new host. Genetic information transfer could be modulated by repressors and activators, which can induced via small molecules or peptides, or in response to excision from the host genome. An intriguing mechanism for blocking DNA transfer between two related donor cells is entry exclusion with many conjugative systems encoding Eex genes as well as associated genes for surface exclusion, which block cell-to-cell contact. It is noteworthy that bacteria such as *Enterococcus* use pheromones to trigger gene expression prior to conjugative DNA transfer with the pheromone being released by the recipient cell. Therefore both temperature and pheromones could be used in a device to modify the conjugation rate.

### Communication and Turing machines

Let us now turn into the relation between computation and metabolism inspired by Turing [11]. Turing states that an organism, most of the time, develops from one pattern into another. Many years later, Bray [12] argued that a single protein is able to transform one or multiple input signals into an output signal, thus it can be viewed as a computational or information carrying element. Following this line of thought, we provide a framework to show that bacteria could have computational capability and act as molecular machines. This relationship is based on the mapping between the metabolism and a RM (equivalent to a Turing Machine, TM). Specifically, we think the reactions in the bacterium as increment/decrement instructions of the RM, where the RM registers count the number of molecules of each metabolite. This approach highlights the mechanical aspect of a bacterium, which can work backwards and forwards executing reactions through its metabolites.

It is well known that a von Neumann architecture is composed of a processing unit, a control unit, a memory to store both data and instructions, and input-output mechanisms. We propose an effective formalism to map the von Neumann architecture to an entire bacterial cell, which becomes a molecular machine. We model the processing unit of the bacterium as the collection of all its chemical reactions, so as to associate the chemical reaction network of bacteria with a TM [13]. Here we use GDMO [5] to obtain Pareto fronts representing multiobjective optimisations in the metabolism. Each point of the Pareto front provided by GDMO is a molecular machine to execute a particular task. Pareto optimality allows to obtain not only a wide range of Pareto optimal solutions, but also the *best trade-off design*. In Figure 3 we show a Pareto front obtained with GDMO when optimising acetate and succinate.

**Fig. 3.**
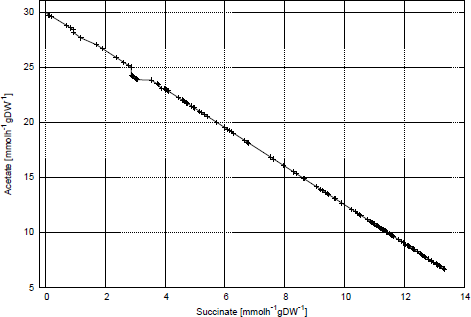
*E. coli* Pareto fronts for the simultaneous maximisation of succinate and acetate production obtained by GDMO in anaerobic conditions (*O*_2_ = 10 *mmolh*^−1^ *gDW*^−1^), with glucose feed equal to 10 *mmolh*^−^ *gDW*^−1^. The acetate represents sources of energy; the succinate enters the krebs cycle.

Optimal genetic interventions in bacteria, framed as optimal programs to be run in a molecular machine, can be exploited to extend and modify the behaviour of bacteria. For instance, programs can instruct cells to make logic decisions according to environmental factors, current cell state, or a specific user-imposed aim, with reliable and reproducible results.

The LGT can be thought of as a process for increasing and decreasing the computational capability of an organism, while seeking the trade-off between computation and minimal genome length. Let *y* be an array representing the sequence of the *L* genes of the organism. During the evolution process, a gene or a subsequence of genes (e.g. an operon) can be duplicated and inserted in the sequence. Without loss of generality, let us assume that the last *k* genes are duplicated:

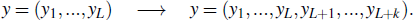

This process is called *gene amplification* or *gene duplication*. It has been estimated that 50% of *E. coli* genes are paralogs i.e. have arisen from a gene duplication event, as opposed to orthologs, which have arisen due to species divergence. The *condition of duplication* holds at the beginning: *y*_*l*_ = *y*_*l*−*k*_, ∀*l* = *L* + 1, …, *L* + *k*, but it is not guaranteed at later steps, due to the fact that the duplication is a stochastic process. Then, after mutations occurring on new and existing genes, the array of genes can be denoted as *y* = (*y*_1_, …, *y*_*L'*_), where *L'* = *L* + *k*. Let us suppose that *y*_*L*_ is responsible for the reaction *D*_*i*_ → *D*_*j*_ + *H*_*r*_ and for the corresponding instruction *inc*(*i*, *r*, *j*) in the Minsky’s register machine (RM, formally defined in Section II) [14], [15]. After the duplication event, also *y*_*L*+*k*_ will be responsible for the same reaction, while after the mutation *y*_*L*+*k*_ will code for another reaction, say *D*_*i'*_ → *D*_*j'*_ + *H*_*r'*_.

The computational complexity of an organism evolves on the basis of stochastic processes and natural selection. Indeed, the mutation process is stochastic, and generates the possibility of adding new instructions to the machine. The natural selection can keep or discard each new instruction. If the new reaction *inc*(*i'*, *r'*, *j'*) is included in the RM, its computational capability increases. We can deduce that each duplication followed by a mutation shapes the computational capability of the metabolic machine represented by its metabolism. These processes permit to increase the range of chemical reactions available in the organism, increasing also the range of increment and decrement instructions in the RM representing its metabolism.

### Metabolic networks as vehicles for communication

Inspired by Brent and Bruck [16], who studied similarities and differences between biological systems and von Neumann computers, we propose a correspondence between the von Neumann architecture and bacteria. Specifically, the metabolism of a bacterium can be viewed as a Turing Machine. The bacterium takes as input the substrates required for its growth and, thanks to its chemical reaction network, produces desired metabolites as output. The string *y* acts as a program stored in the RAM [13]. Let us consider the multiset *Y* of the bits of *y*. A partition П of the multiset *Y* = {*y*_1_, *y*_2_, …,*y*_*L*_} is a collection {*b*_1_, *b*_2_, …, *b*_*p*_} of submultisets of *Y* that are nonempty, disjoint, and whose union equals *Y*. The elements {*b*_*s*_}_*s*=1_, …, _*p*_ of a partition are called blocks. We denote by *P*(*Y*; *p*) the set of all partitions of *Y* with *p* blocks. *P*(*Y*; *p*) has a cardinality equal to the Stirling number, namely ∣*P*(*Y*; *p*)∣ = *S*_*L*,*p*_.

In order to formalise the control unit behaviour, let us define the function:

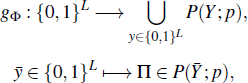

where the partition Π is uniquely determined by the pathway-based clustering of the chemical reaction network. We can formalise this clustering as a *p*-blocks partition Φ of the set of the bit indexes in the string *y*. In particular, if we denote by [*L*] the set of the first *L* natural numbers, we have Φ ∈ *P*([*L*]; *p*) [13]. The partition Φ allows the control function *g*_Φ_ to partition the multiset *Y* associated with the string *y*. Each element of the partition Π is the submultiset *b*_*s*_ of all the gene sets related to reactions in the *s*th pathway. The processing unit of the bacterium could be modelled as the collection of all its chemical reactions. Therefore, the chemical reaction network of bacteria can be associated with a TM [17]. Let us consider the *Minsky’s register machine*, i.e. a finite state machine augmented with a finite number of registers. Formally, a Minsky machine *ℳ* = (*D*, *i*_0_, *i*_1_, φ) is composed of a finite set *D* of states, a finite set *H* = {*H*_*r*_}_*r*_ of registers, and a multivalued mapping φ : *D*\{*i*_0_} → {(*H*_*r*_, *i*), (*H*_*r*_, *j*, *k*) ∣ *H*_*r*_ ∈ *H*, *j*, *k* ∈ *D*}. The set *D* has two distinguished elements *i*_0_, *i*_1_ ∈ *D* representing the initial state and the halting state respectively. Each register *H*_*r*_ of the RM stores a non-negative integer. The instruction *inc*(*i*, *r*, *j*) increments register *r* by 1 and causes the machine to move from state *i* to state *j* through the mapping φ(*i*) = *j*. Conversely, the instruction *dec*(*i*, *r*, *j*, *k*), given that *H*_*r*_ > 0, decrements register *r* by 1 and causes the machine to move from state *i* to state *j* (φ(*i*) = *j*); if *H*_*r*_ = 0, the machine moves from state *i* to state *k* (φ(*i*) = *k*). The Minsky’s RM has been proven to be equivalent to the TM ^1^. Indeed, a RM is a multitape TM with the tapes restricted to act like simple registers (i.e. “counters”). A register is represented by a left-handed tape that can hold only positive integers by writing stacks of marks on the tape; a blank tape represents the count ‘0’. The chemical reaction network of a bacterium can be mapped to the RM by defining [17]: (i) the set of state species {*D*_*i*_}, where each *D*_*i*_ is associated with the state *i* of the RM; (ii) the set of register species {*H*_*r*_}, where each *H*_*r*_ is associated with the register *r* of the RM, and therefore represents the molecular count of species *r*. The instruction *inc*(*i*, *r*, *j*) represents the chemical reaction *D*_*i*_ → *D*_*j*_ + *H*_*r*_, while the instruction *dec*(*i*, *r*, *j*, *k*) represents either *D*_*i*_ + *H*_*r*_ → *D*_*j*_ or *D*_*i*_ → *D*_*k*_ depending on whether *H*_*r*_ > 0 or *H*_*r*_ = 0 respectively. The molecular machine performs the “test for zero” by executing the reaction *D*_*i*_ → *D*_*k*_ only when *H*_*r*_ is over, since the *r*th register cannot be decreased and the reaction *D*_*i*_ + *H*_*r*_ → *D*_*j*_ cannot take place. In the FBA approach coupled with the metabolic machine, the variables are the fluxes of the chemical reactions, therefore a high flux corresponds to both a high rate of reaction and a high mass of products. Hence, given the increment reaction *inc*(*i*, *r*, *j*), the value of *H*_*r*_ is positively correlated with the reaction flux; conversely, in the decrement reaction *dec*(*i*, *r*, *j*, *k*), when *H*_*r*_ > 0 the value of *H*_*r*_ is negatively correlated with the reaction flux. In a fixed volume *V* in which the reactions occur, given two reactions *inc* and *dec* with fluxes *v*_1_ and *v*_2_ respectively, the metabolism of the bacterium has a probability of error per step equal to ε = *v*_2_/(*v*_1_/*V* + *v*_2_).

Since the simulated TM can be universal, the correspondence between metabolism and TM allows to perform any kind of computation through a set of species and chemical reactions characterised by their flux. As a result, bacteria can carry out at least any computation performed by a computer. A program embedded in a bacterium, whose metabolism works like a TM, could be able to implement the robust knockout strategy found by GDMO [5].

It is noteworthy that, in theory, the frequency and species restrictions of future gene lateral transfer capacity could be encoded in the genes being transferred to a bacterium. This could result into changing the size of the mobilome. The metabolic computing could also generate different partition of bacteria that could share genetic material and co-evolve accordingly.

## III. Geobacter: network communication through biofilms

An anaerobic bacterium, *Geobacter* found in sediment under the Potomac River in 1987 has soil bioremediation capacity due to its ability to respire iron and other metals. In 2002 microbiologists further discovered that it could produce electricity from the organic matter found in soils, sediments and wastewater due to electrically conductive pili, sort of hairs [18], [19], [20]. These conductive protein appendages that transfer electrons to metal oxides and to other cells are only 3 to 5 nanometers in diameter and more than a thousand times longer than they are wide. The pili of a population of this bacterium form an electro-active biofilm [21] that provides a direct electrochemical connection with the electrode surface using it as electron exchanger, without the aid of mediators. *Geobacter* species produce higher current densities than any other known organism in microbial fuel cells (for example Rhodoferax ferrireducens, Shewanella [22]) and are common colonisers of electrodes harvesting electricity from organic wastes and aquatic sediments. On electrodes, the bacteria produce thick, electrically conductive biofilms. These processes can be harnessed for the bioremediation of toxic metals and the generation of electricity in bioelectrochemical cells. Key to these applications is a detailed understanding of how these nanostructures conduct electrons. This discovery not only represents a paradigm shift in biology but also in nanotechnology.

### Useful Geobacter properties for communication applications

Our studies of Pareto fronts highlight the potentialities of choosing different strains according to the biomass - electronic properties. A larger biomass may turn into more dense biofilms, while optimising in a different way could allow to obtain better electronic properties of the biofilm. *Geobacter* has been shown to produce biofilms containing exopolysaccharides [23] as well as proteinaceous structures (pili). The biofilm matrix or extracellular polymeric substance (EPS) or xap (extracellular anchoring polysaccharide) is a perfect medium for entrapment of redox proteins for short-and long-range electron transfer, in particular localisation of essential cytochromes beyond the *Geobacter* outer membrane.

The biofilm, i.e. a cohesive aggregate of billions of cells, can conduct electrons. Specifically, networks of microbial nanowires that are able to conduct electrons, course through the biofilm and can move charges over significant distances, e.g. thousands of times the bacterium’s length. Since these protein filaments can conduct electrons along their length, the biofilm is turned into a sort of a metal that can conduct electrons as far as the biofilm can be extended. The problem of electrical isolation of a group of redox-active organisms can be associated with the problem often found in human cities. Namely, the electrical isolation can cause local variations in potential, thus resulting in damage to individuals and potentially to the whole group of organisms. Therefore, the group can benefit from a link to a common ground [24]. The redox-active enzymes can be spatially distributed on a biofilm, which provides a structural matrix for enzymes, metal and mineral substrates.

Cultivation and analysis of individual bacterial species has been at the core of experimental microbiology for more than a century. Microbial communities rather than individual species generally control process rates and drive key biogeochemical cycles. Recently a community of several bacteria has been investigated. cellobiose served as the carbon and energy source for C. cellulolyticum, whereas *D. vulgaris* and *G. sulfurreducens* derived carbon and energy from the metabolic products of cellobiose fermentation and were provided with sulfate and fumarate respectively as electron acceptors [25]. Recent experiments have shown that *Geobacter*’s electric nanowire has still large evolutionary potential, which we believe can be explored using optimisation algorithms. Lovley was able to evolve a new strain that dramatically increases power output per cell and overall bulk power [26]. The concentrations needed on the electrode to produce electricity are reached faster, due to the biofilm being thinner than earlier strains. In this experiments, he added a tiny pushback current in the electrode in order to select those *Geobacter* species able to press harder to get rid of the electrons given by the additional current. As a result, with respect to the original strain, he obtained the evolution of a microorganism able to press at least eight times more electric current across the electrode [27].

### Conclusions

In this paper we have highlighted the links between bacterium communication, gene duplication, lateral gene transfer events and metabolic complexity. The methodology we propose allows to optimise simultaneously several objectives, i.e. the output of the metabolic “computation” versus communication carried out by bacteria. Population effects (a group of same cells) and community effects (a group of different cells) can be also investigated with optimisation techniques.

A Pareto front can help investigate the trade-off between computation and communication. This is equivalent to identify the optimal size of the “communication” part of the metabolic network, namely the set of reactions responsible for movement aimed at seeking food. For instance, while the whole metabolic network of the *Geobacter* is responsible for the bacterial computation, the subset of reactions responsible for communication is involved in the production of biofilm. This approach highlights the complex behaviour that may arise in molecular machines; although nano communication networks and synthetic biology are still in their infancy, we foresee the potentialities to build and optimise synthetic organisms that could be designed for optimising specific communication performances and networks.

In groups of multiple interacting microbial populations, also called *microbial consortia*, the recent work by Ji et al. [28] showed that when the consortium of bacteria is thought of as consisting of logic operating cells, it can compute Boolean functions. In a group of interacting bacteria, the balance of the intra-cellular amount of energy depends on the properties of the genes and their codon usage, whereas the global production of energy may depend on the topology of the network [29]. These recent findings, if coupled with our work, lead to speculate that colonies of bacteria can compute both the optimal intra-cellular configuration for energy production and the optimal design for the energy production of the whole consortium.

## Footnotes

1 M.L. Minsky. Computation. Prentice-Hall, 1967.

